# Modeling disease spread in populations with birth, death, and concurrency

**DOI:** 10.1101/087213

**Authors:** Joel C. Miller, Anja C. Slim

## Abstract

The existence of sexual partnerships that overlap in time (concurrent relationships) is believed by some to be a significant contributing factor to the spread of HIV, although this is controversial. We derive an analytic model which allows us to investigate and compare disease spread in populations with and without concurrency. We can identify regions of parameter space in which its impact is negligible, and other regions in which it plays a major role. We also see that the impact of concurrency on the initial growth phase can be much larger than its impact on the equilibrium size. We see that the effect of concurrency saturates, which leads to the perhaps surprising conclusion that interventions targeting concurrency may be most effective in populations with low to moderate levels of concurrency.

**Author Summary:** We consider the spread of an infectious disease through a population modeled by a dynamic network with demographic turnover. We develop a stochastic model of the disease and derive governing equations that exactly predict the large population (deterministic) limit of the stochastic model. We use this to investigate the role of concurrency and find that interventions targeting concurrency may be most effective in populations with lower levels of concurrency.

Our model is not intended to be an accurate representation of any single population. Rather it is intended to give general insights for intervention design and to provide a framework which can be further specialized to particular populations.

This model is the first model to allow for analytic investigation of the impact of concurrent partnerships in a population exhibiting demographic turnover. Thus it will be useful for investigating the “concurrency hypothesis.”

## Introduction

The HIV epidemic has had significant impact worldwide, but it has had an especially large impact in SubSaharan Africa [1]. The reasons for this difference are many, complex, and not fully understood. One proposed factor is a larger frequency of sexual partnerships that overlap in time, the so-called “concurrency hypothesis” [2, 3]. The concurrency hypothesis has received significant attention, but it is controversial [4–10].

The development and evaluation of potential interventions against infectious disease spread requires us to be able to predict the spread of the disease with and without the intervention in place. A crucial factor that must be considered in evaluating any potential intervention is the indirect effects of the intervention: If we directly preventing one infection from occurring, then this also prevents that individual from transmitting onwards, thus indirectly reducing exposures of other individuals. In turn their reduction in exposure reduces potential exposures of others further down the line.

Such a feedback mechanism is an inherently nonlinear effect, and so the full outcome of an intervention will depend nonlinearly on the direct effect. Doubling the intervention effort will not double the intervention impact. It may have greater or lesser impact depending on the magnitude of the indirect effect. Because of this, we often rely on mathematical models to predict the total intervention outcome. The required model complexity depends on the intervention under consideration, the biological properties of the disease, and the contact structure of the population.

Most mathematical models of disease spread ignore the fact that partnerships have duration. This is reasonable in a population with many partnerships and low transmission probabilities [11], but for a disease such as HIV, the relevant partnerships are often long-lasting and the typical number of overlapping partnerships is generally small. To be relevant for the concurrency hypothesis, we require that the mathematical model incorporate partnership duration in some form. Without it, it is impossible to consider partnerships that overlap in time. Additionally, given the time-scale of the HIV epidemic, demographic turnover plays an important role. There has not been a mathematical model which captures all of these features, and so most investigations turn to simulation, many of which are reviewed in [12].

The absence of an analytic model contributes to a large gap in our ability to understand concurrency. Analytic models make it much easier to interpret model predictions, to specifically isolate the impact of different effects, and to clarify underlying assumptions. Simulated results may be difficult to replicate by other researchers, the slow process of simulation may make parameter space impossible to investigate, and it may be difficult to identify which assumptions of the simulation are responsible for which outcomes. All of this contributes to the ongoing controversy about concurrency, and even miscommunication within the field. With an analytic model, these issues are significantly reduced.

Recent work [13–17] has shown that it is possible to make mathematical models for SI and SIR disease spread in networks with only a handful of equations. The approach has been generalized to networks with changing partnerships, but still assume a closed population. Because the HIV epidemic developed over decades, it is not appropriate to ignore individuals entering and leaving the at risk-population as they age.

In this paper we adapt the approach of [15–17] in order to accommodate “births” and “deaths” representing entry into and exit from the at-risk population. We show that the resulting equations accurately predict the outcome of simulations in large populations, and our primary focus is on using the model to investigate the role that concurrency can play in the spread of a “Susceptible–Infected” (SI) disease such as HIV.

### Concurrency

A potentially important, but incompletely understood, factor in the spread of HIV and other sexually transmitted diseases is the role of “concurrent” relationships as compared to serial monogamy [2, 3]. Whether its impact is significant is hotly debated [4–10].

To highlight the potential role of concurrency, we consider an individual “Alex” who has two partners “Bobbie” and “Charlie” over a period of 1 year. We demonstrate two potential partnership arrangements and explore their impact on disease transmission in Fig. 1.

**Fig 1.**
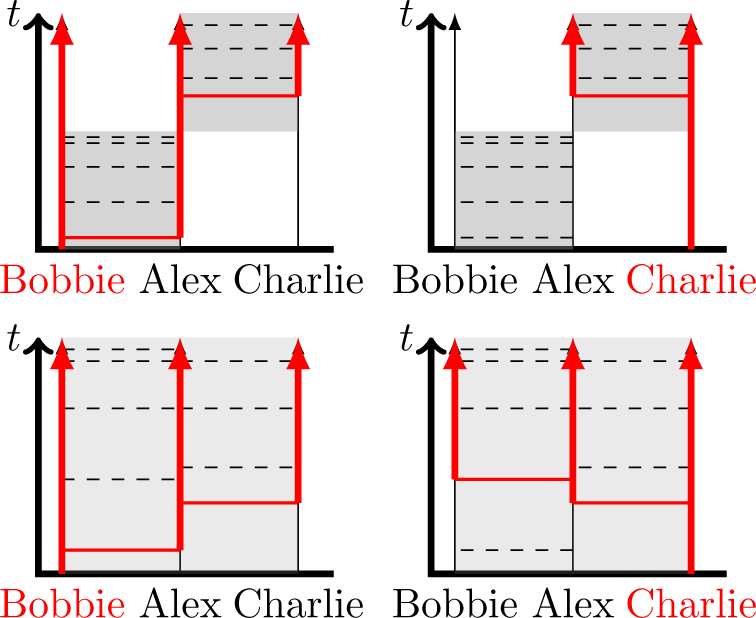
Sample scenarios comparing concurrent and serially monogamous relationships: Shaded regions denote the existence of a partnership. Dashed lines denote infection opportunities within the relationship that would cause infection if one individual were susceptible and the other infected. Vertical red lines denote time in which an individual is infected, and horizontal red lines denote successful transmissions. In the concurrent case, we keep exactly the same number of transmission opportunities with the gaps exactly doubled, assuming that the interaction rate within each partnership is half that of the serial case. Concurrency can speed up onward transmission and provide additional transmission routes.

In the first scenario, the partnership with Bobbie lasts for six months and is replaced by a six month partnership with Charlie (the serial case), while in the second scenario both partnerships overlap for the entire period (the concurrent case). In the concurrent case, the intervals between transmitting events are doubled.

Let us focus first on Alex’s risk of infection if Bobbie is initially infected. In both cases, Alex has the same probability of becoming infected, though infection would occur sooner in the serial case. In contrast, if Charlie is initially infected, Alex would become infected sooner in the concurrent case (but again with the same overall probability). So the two models appear to give similar outcomes from the point of view of the cumulative risk of infection to Alex.

For onward transmission, however, we see the potential for large differences. If Bobbie is initially infected, transmission from Bobbie to Alex to Charlie will tend to happen faster in the concurrent case because there is no built-in delay while waiting for the partners to change.^1^ In turn this allows Charlie to begin transmitting to other partners earlier. In addition, if Charlie is initially infected, transmission from Charlie to Alex to Bobbie is possible only in the concurrent case.

It is important to recognize that the “concurrency hypothesis” does not suggest that there is a difference in the total risk of infection of an individual who has multiple concurrent partners compared to one who has the same number of partners sequentially. Instead, it increases the risk to the partners.

As a general rule we anticipate that concurrency will increase the spread of disease through two mechanisms:

1. by allowing the disease to trace transmission routes faster, and
2. by providing additional transmission routes.

In the low transmission rate limit, for which the possibility of transmitting multiple times is negligible, we can assume that each partnership either transmits once or not at all. In this case, Fig. 2 suggests that the difference between concurrent and serial relationships is likely to be insignificant.

**Fig 2.**
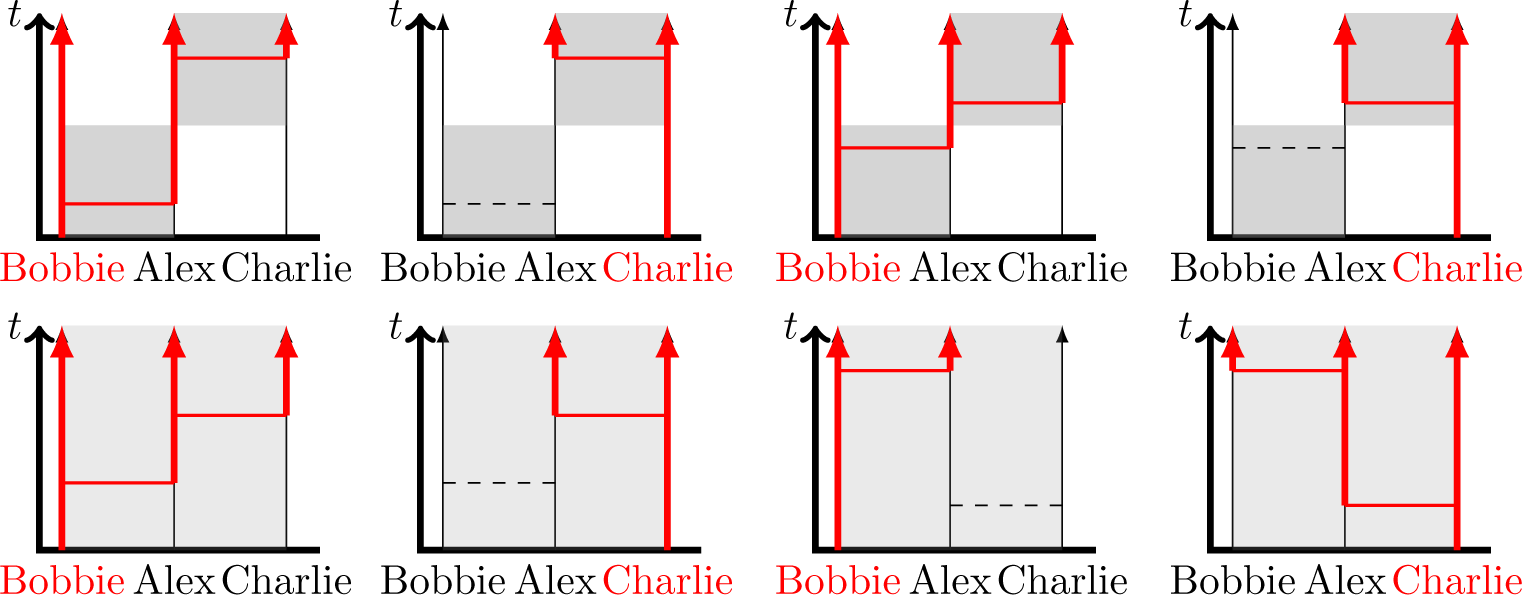
Scenarios for low transmission rates: In the low transmission rate limit where we can assume at most one transmission happens per partnership, the additional transmission paths and faster transmissions from concurrent partnerships do not play a significant role. There are two cases of interest: The first two columns show the case in which the Bobbie-Alex partnership transmits relatively earlier in the partnership than the Alex-Charlie partnership. The last two columns show the Alex–Charlie partnership transmitting faster.

Conceptually we can explain this by imagining that we only observe transmission events. Anything that influences the disease spread must be detectable by our observations. Conversely, if we cannot detect something by observing disease transmission events, then it cannot matter to the disease spread. In particular, if no partnerships transmit more than once, then we cannot tell whether an individual who is transmitting is doing so to one of several randomly selected current partners or if it is transmitting to its current partner who is randomly selected from the potential partners.

## Materials and Methods

Our goal is twofold:

1. To demonstrate an analytic model for disease spread in a dynamic population with concurrency and show that it accurately predicts simulations in large populations.
2. To use the model developed to provide guidance about the role of concurrent relationships in simplified scenarios.

In this section we introduce our stochastic population and disease model, state the governing equations for the large population limit (the equations themselves are derived in the Supporting Information), and demonstrate that they accurately reproduce simulated epidemics in large populations.

Part of the reason for the debate about concurrency’s effect can be traced to the large parameter space and the amount of time it takes to simulate for different parameters. Simulations have studied only a relatively small portion of parameter space, and so some of the disagreement revolves around intuition about how parameter changes would affect outcomes rather than quantitative statements. Our analytic model will allow for a much more efficient exploration of parameter space.

### Population/Disease Assumptions

We will use a discrete-time model. Each time step is broken down into sub-intervals in which the actions occur in a specific order, shown in Fig. 3. At the start of a time step, each partnership connecting a susceptible and infected individual transmits with probability *τ*. Then some individuals leave the population, each independently with probability **μ**. Next *μN* new individuals arrive where *N* is the typical population size. Then each remaining partnership ends independently with probability *η*. Finally new partnerships are formed until all individuals reach their target number of partners. These assumptions are similar to those of [18, 19].

**Fig 3.**
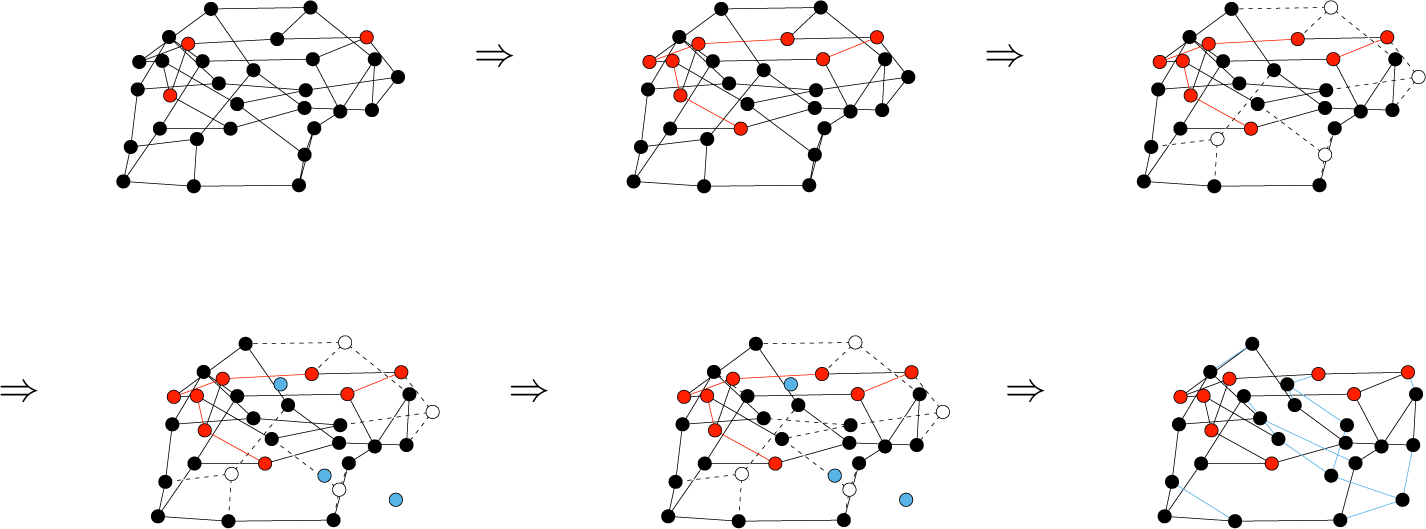
Sequence of events in each time step. We begin with a network with some infected individuals (red). Then infected individuals transmit to some partners (red edges). Then some individuals leave the population (white). Other individuals are born (blue). Then some edges break (dashed). Finally edges are added so that the new individuals, the individuals whose partners left, and the individuals whose edges broke all return to their target number of partners. The sequence then repeats.

We will present an analytic model that captures the deterministic limit of these assumptions. We have chosen a discrete-time model because the complementary simulations are easier in a discrete-time framework. In a large enough simulated population with a large enough time step, many partnerships end and many new individual enter at each time step. Thus an individual who needs a new partner has many choices. In a continuous-time model or with a small time step (relative to population size), it would be more difficult for an individual who has recently ended a partnership to find a new partner.

For simulations, we choose the time step of the discrete-time framework to balance competing interests. We want a small time step so that *μ*, *η*, and *τ* are small (at leading order they are proportional to the time step). When they are not small, the somewhat arbitrary order of events we have assumed will begin to impact outcomes. However, too small of a time step will mean that few partnerships end in a time step. This makes it difficult for nodes to immediately find new partners. As the population size increases this becomes less of a problem so smaller time-steps become feasible. However, the computational effort becomes greater. This is not an issue in the analytic model, and so a continuous-time model is not difficult to consider. The predictions from the continuous-time and discrete-time models will be the same for small parameters.

### Governing equations

We derive the governing equations in the Supporting Information. These equations are based on the “Edge-based Compartmental Modeling” approach of [13, 15, 16]. They are low-dimensional, but are significantly more involved than even the dynamic network models presented in [16]. This is because the age of an individual gives us some information about that individual’s status. In addition the age of a partnership gives information about the age of the partner. Thus in calculating the risk an individual has from its partners, we need to account for the probability the partner has a given status, which depends on the age of the partner, which in turn depends on the age of the edge, which itself is dependent on the age of the individual. To sort out the dependencies, age of individual and age of partnership are needed as independent variables.

Table 1 defines the variables and parameters of the model.

**Table 1.** The variables for our simulations and equations.

| Variable | Description |
| --- | --- |
| $t$ | Time |
| $u$ | The test individual |
| $a_u$ | The age of test individual $u$, measured so that $a_u = 0$ in the first time step after $u$ is born. |
| $a_e$ | Age of an edge of interest, measured so that $a_e = 0$ in the first time step after the edge forms. |
| $S(t)$ | The proportion of the population that is susceptible at time $t$, equivalently the probability a randomly selected individual is susceptible at time $t$, or equivalently the probability a test individual is susceptible at time $t$. |
| $I(t)$ | $1 - S(t)$: The proportion infected at time $t$. |
| $s(t, a_u)$ | The probability a test individual of age $a_u$ is susceptible at time $t$. |
| $\Theta(t, a_u)$ | The probability a stub belonging to $u$ has not transmitted infection to it from a partner by the start of time step $t$. |
| $\Phi_S(t, a_u)$ | As for $\Theta$, but with the additional requirement that at the start of time step $t$ the partner is susceptible. |
| $\Phi_I(t, a_u)$ | As for $\Theta$ (no partner has transmitted to $u$ through the stub), but with the additional requirement that at the start of time step $t$ the partner is infected. |
| $\phi_S(t, a_u, a_e)$ | The probability that a stub belonging to an age $a_u$ individual has not transmitted infection to it by time $t$, is connected to a susceptible neighbor, and the current partnership has age $a_e$. |
| $\chi(t, a_e)$ | The probability that an age $a_e$ edge of a test individual connects to a susceptible individual. |
| $\psi(x)$ | $\sum_{k=0}^{\infty} P(k)x^k$: The probability generating function of the degree distribution. |
| $\mu$ | The probability a random individual will leave the population in a given time step. |
| $\tau$ | The transmission probability in a time step. |
| $\eta$ | The probability a partnership will end in a time step. |
| $p_b$ | $1 - (1 - \mu)(1 - \eta)$: The probability that a test individual's partnership will end (break) either because the partnership ends naturally $\eta$, or the partner leaves the population $\mu$. It does not include the possibility of the test individual leaving. |
| $P_e$ | $\frac{(1-\mu)(\eta+\mu-\eta\mu)}{(1-\mu)(\eta+\mu-\eta\mu)+\mu}$: The probability that a newly formed partnership will be with a previously existing individual. |
| $\rho$ | The proportion of the population randomly infected at $t = 0$. |
| $b$ | The number of individuals entering the population each time step (assumed to be constant). |
| $N$ | $b/\mu$: The average population size. |

The model parameters are the death probability *μ*, the transmission probability *τ*, the partnership change probability *η*, and the initial fraction infected *ρ*. We seek the susceptible and infected fractions of the population *S* and *I*. The governing equations are

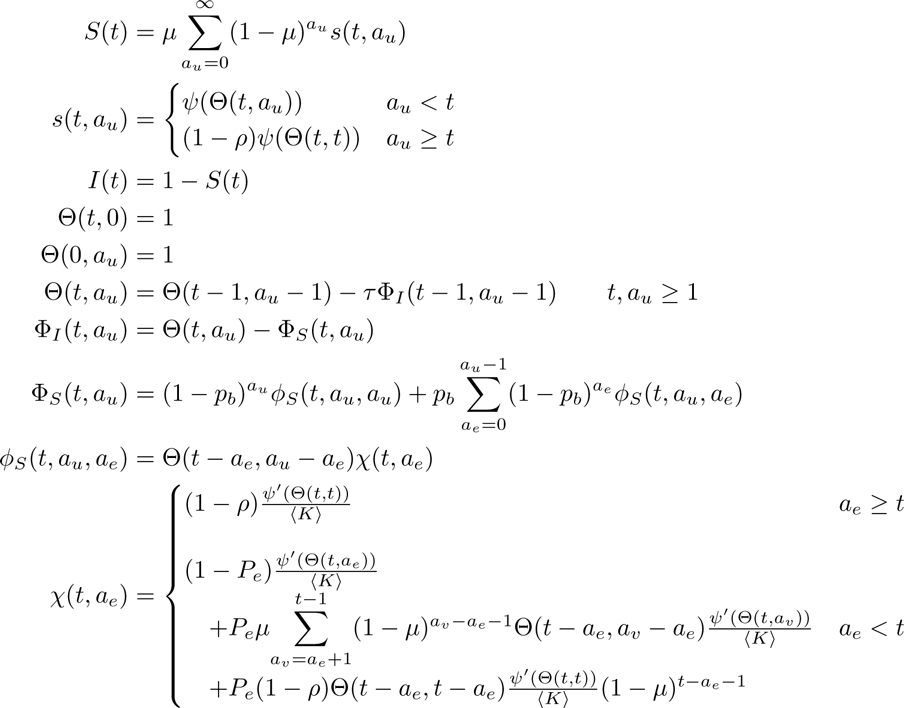

where *p_b_* = 1 − (1 − *η*)(1 − *μ*) is the probability an individual’s partnership ends in a time step either because of partnership change or the partner leaving the population, and *ψ*(*x*) = ∑_*k*_*P*(*k*)*x^k^* is the probability generating function of the degree distribution. We derive this model and a continuous-time differential equations version in the Supporting Information.

### Comparison with simulation

To demonstrate the accuracy of the equations and provide some insight into model predictions, we compare predicted dynamics (solutions of the system of equations) with stochastically simulated epidemics under different conditions in Figs. 4 and 5.

**Fig 4.**
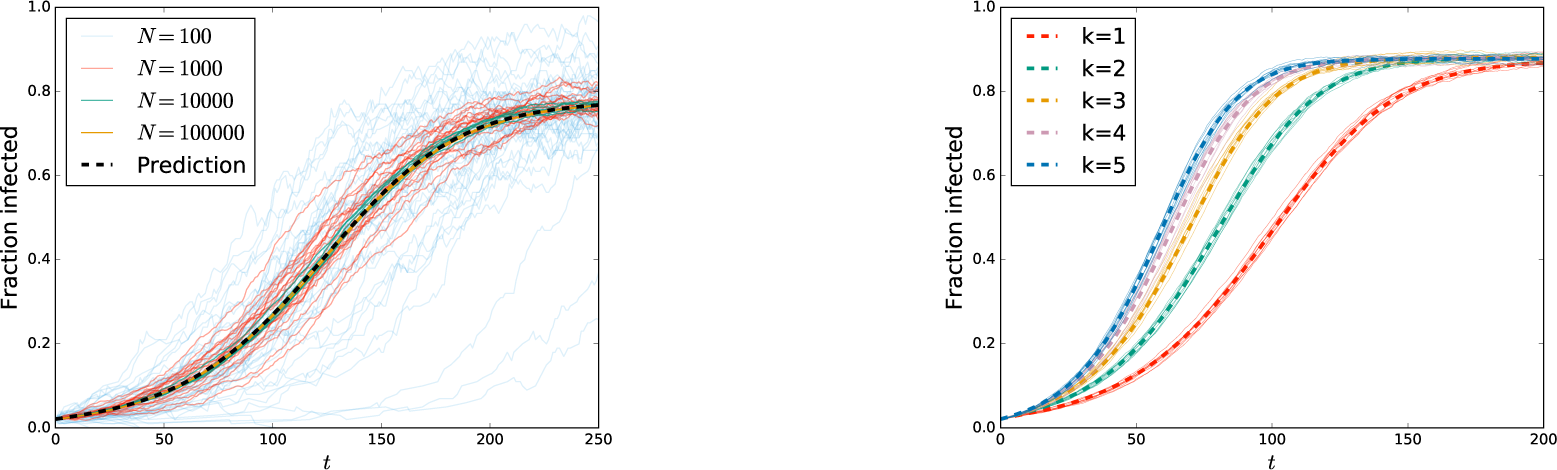
Comparison of dynamics from analytic prediction and stochastic simulation: (Left) We consider stochastic epidemic simulation in populations of average size *N* = 102, 103, 104 and 105 with *κ* = 3, *η* = 0.2/3, *τ* = 0.05/3, *ρ* = 0.02, and *μ* = 0.01. As *N* increases, the simulations converge to the prediction. (Right) For *η*_1_ = 0.1, *τ*_1_ = 0.1, *ρ* = 0.02, *μ* = 0.01 and average population size *N* = 10000 we compare predictions and simulations for different values of *k*, using *τ* = *τ*1*/k* and *η* = *η*_1_/*k*. We find excellent agreement.

We start Fig. 4 with a plot showing that as the population size increases, the simulations converge to the predicted dynamics for one set of parameters.

The second plot of Fig. 4 considers populations in which all individuals have *k* partners for *k* = 1, 2, 3, 4, and 5 with *τ* = *τ*_1_/*k* and *η* = *η*_1_/*k* for various values of *τ*_1_ and *β*_1_. Simulations and predictions are a good match for different values of *k*. Interestingly we see that for different values of *k* the equilibrium level does not vary much, but the early growth rate does.

None of the populations in Fig. 4 have heterogeneous degree. Figure 5 looks at disease spread in populations with heterogeneous degrees, again showing excellent agreement between stochastic simulation and predictions.

**Fig 5.**
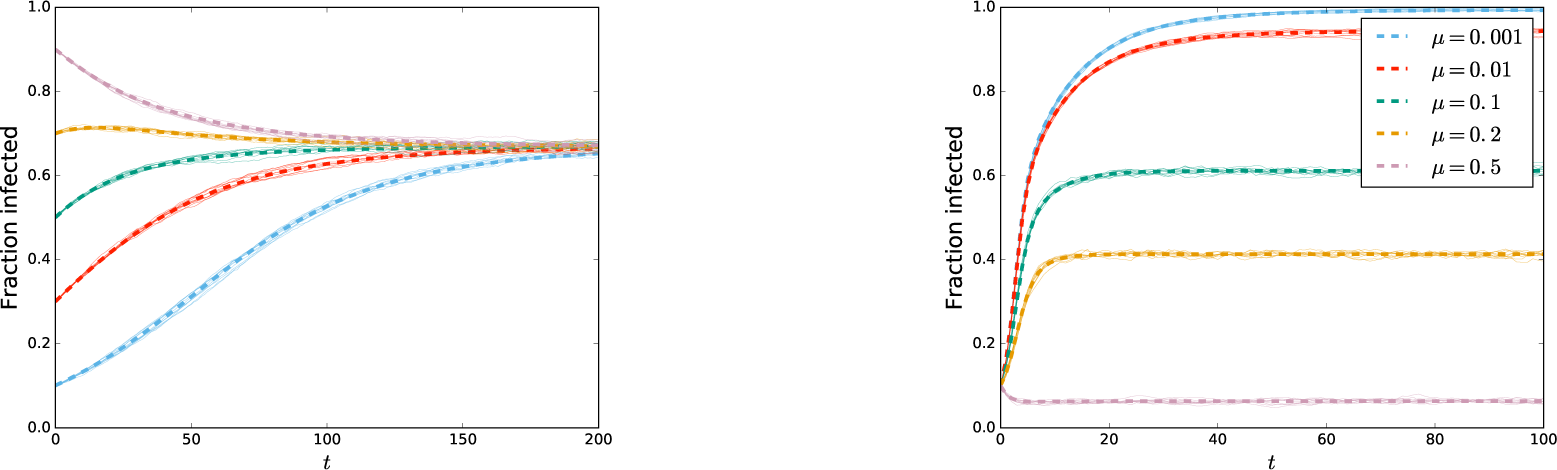
Comparison of dynamics from analytic prediction and stochastic simulation in populations with heterogeneous degree: (Left) Disease spread in populations with average size *N* = 10000 and degree probabilities *P* (2) = *P* (7) = 1*/*2. The parameters are *τ* = 0.01, *η* = 0.005, and *μ* = 0.01. The initial fraction infected varies. (Right) Disease spread in populations with average size *N* = 10000 and degree probabilities *P* (1) = 1*/*2, *P* (10) = 1*/*3, and *P* (20) = 1*/*6. The parameter *μ* varies between populations. The remaining parameters are *η* = 0.05 and *τ* = 0.1. In both plots the dashed curves are predictions and thin solid curves are stochastic simulations.

From the first plot in Fig. 5 we infer that for a given population, the initial proportion infected does not influence the final state. Interestingly, we see that it is possible for the disease to initially grow and then decay for some region of *ρ*. This is because in this region the number of infected high degree nodes increases quickly from the initial condition while the number of infected low degree nodes decreases slowly. At long time, reach a steady state, with the decrease of low degree infections outweighing the increase in high degree infections.

From the second plot of Fig. 5, we infer that increasing the population turnover rate decreases the proportion infected. This is not particularly surprising as it implies that infected individuals leave the population sooner, having had less opportunity to cause further infections.

Our main conclusion from Figs. 4 and 7 is that the equations accurately predict the dynamics of simulations regardless of the parameters used. The numerical solution of these equations is much faster than simulation.

## Results & Discussion

The previous section showed that the analytic equations provide accurate predictions of the disease dynamics in the large-population limit. We now specifically explore the predictions the model makes if we change the amount of concurrency in the population. The second plot of figure 4 hints at our results. We generally see relatively little impact of concurrency on the proportion infected at equilibrium, while we do see an impact on the early growth.

Although the equations allow different individuals to have a different number of partners, for the remainder of this paper we focus on populations in which all individuals have the same number of partners. In particular, this eliminates the need to consider how per-partnership transmission rates may depend on the individual’s number of partners. This allows us to separate the effect of concurrency from the effect of some individuals having more frequent sexual activity than others.

So for our purposes, *ψ*(*x*) = *x^k^*, 〈*K*〉 = *k* for some fixed value *k*. As a base-case, we consider serial-monogamy where each individual has a single partner (*k* = 1). Transmission occurs in a time-step with probability *τ*_1_ and partnerships end with probability *η*_1_. Individuals leave the population with probability *μ*. We compare this with homogeneous populations having concurrency.

We assume that each population is arranged such that the number of partners an individual has over a long period of time is the same. This implies that *η* = *η*_1_/*k* so that the *k* partnerships each lasts *k* times as long. Similarly we assume that the expected number of transmissions an infected individual would cause in a time step is the same. This requires *τ* = *τ*_1_/*k*.

Because of how we have set up our populations, we can rule out many causes for the differences we see. We know that the effects we observe are not explained by within-population heterogeneity in degree, within-population heterogeneity in sexual activity rates, between-population differences in typical life-time number of partners, or between-population differences in the number of transmissions an individual causes per time step. All of these effects have been removed. The only remaining difference between the populations is the number of concurrent partnerships.

### Impact on equilibrium size

We see in Fig. 6 that the impact of concurrency on the equilibrium epidemic size can be significant, but that the effect of increasing *k* saturates quickly. Little changes once *k* > 2. This suggests that reducing concurrency may play an important role in reducing the epidemic size, but that if there is significant clustering, a small or moderate change is unlikely to have a significant impact.

**Fig 6.**
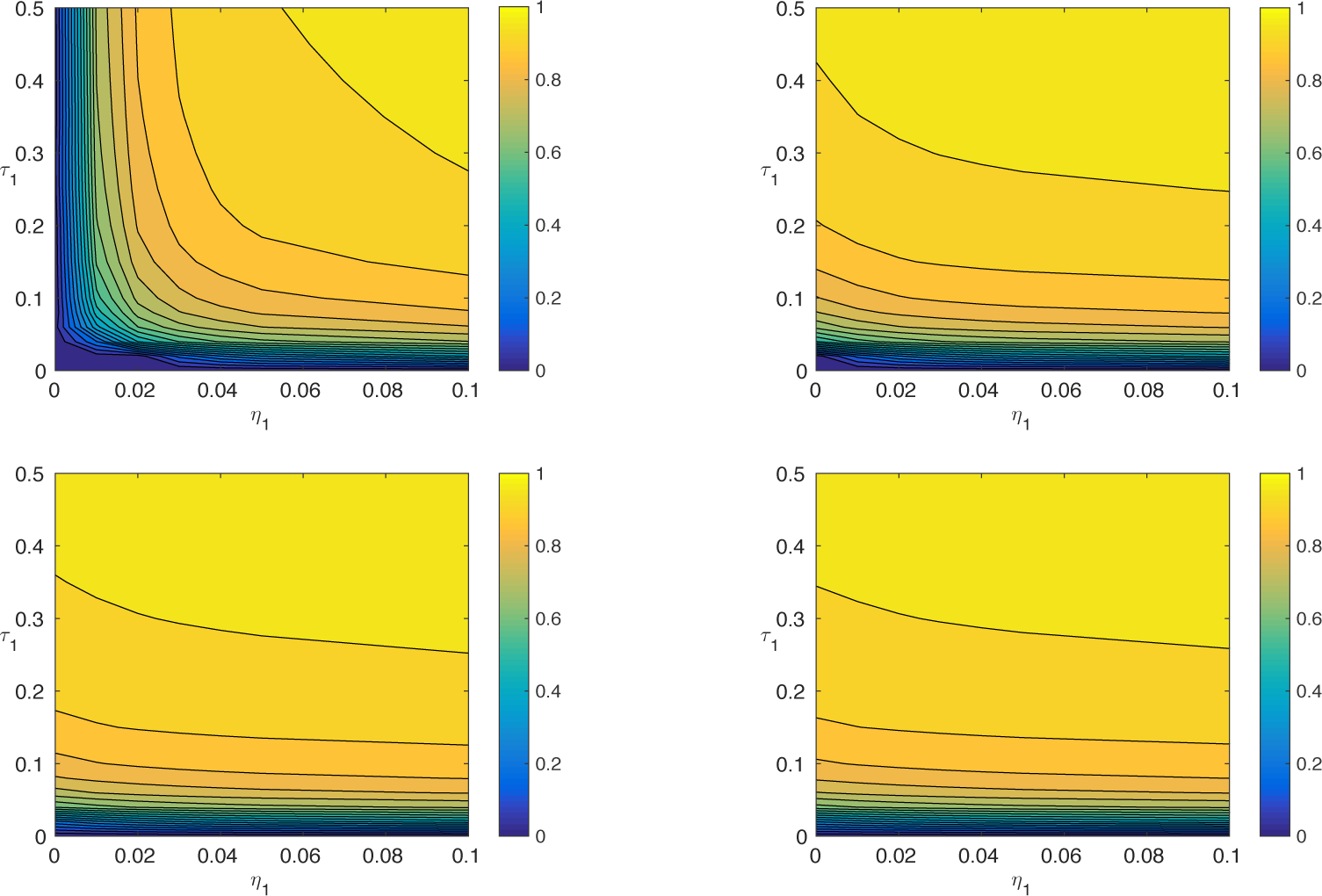
Comparison of equilibrium sizes for different values of *k*: The equilibrium fraction infected for *μ* = 0.01 and different *η*_1_ and *τ*_1_. We use the same axes *η*_1_ and *τ*_1_, taking *η* = *η*_1_/*k* and *τ* = *τ*_1_/*k*. As *k* increases, the figures quickly converge. The effect of concurrency on the equilibrium size quickly saturates.

From the perspective of intervention design, this suggests that interventions which aim to reduce concurrency may be most effective in populations with lower rates of concurrency already. We can think of this analagously with interventions targetting *R*_0_ in an SIR disease. Reducing *R*_0_ from a large number to a moderate number has little effect on the predicted final size, while a reduction from a moderate number to a small number (even if still above 1) would have a much larger impact.

We note that close to the epidemic threshold the impact of any parameter change (including amount of concurrency present) can be significant.

### Impact on early growth

Although Fig. 6 appears to suggest concurrency is unimportant, this is only for the equilibrium size. We saw in Fig. 4 that concurrency can play a role when we consider the early growth rate of the disease even if the equilibrium size is unaffected. That is, concurrency facilitates early spread of a disease. We demonstrate this more generally in Fig. 7. Although the impact of increasing *k* will eventually saturate, it takes much longer before saturating for early growth than for equilibrium size.

**Fig 7.**
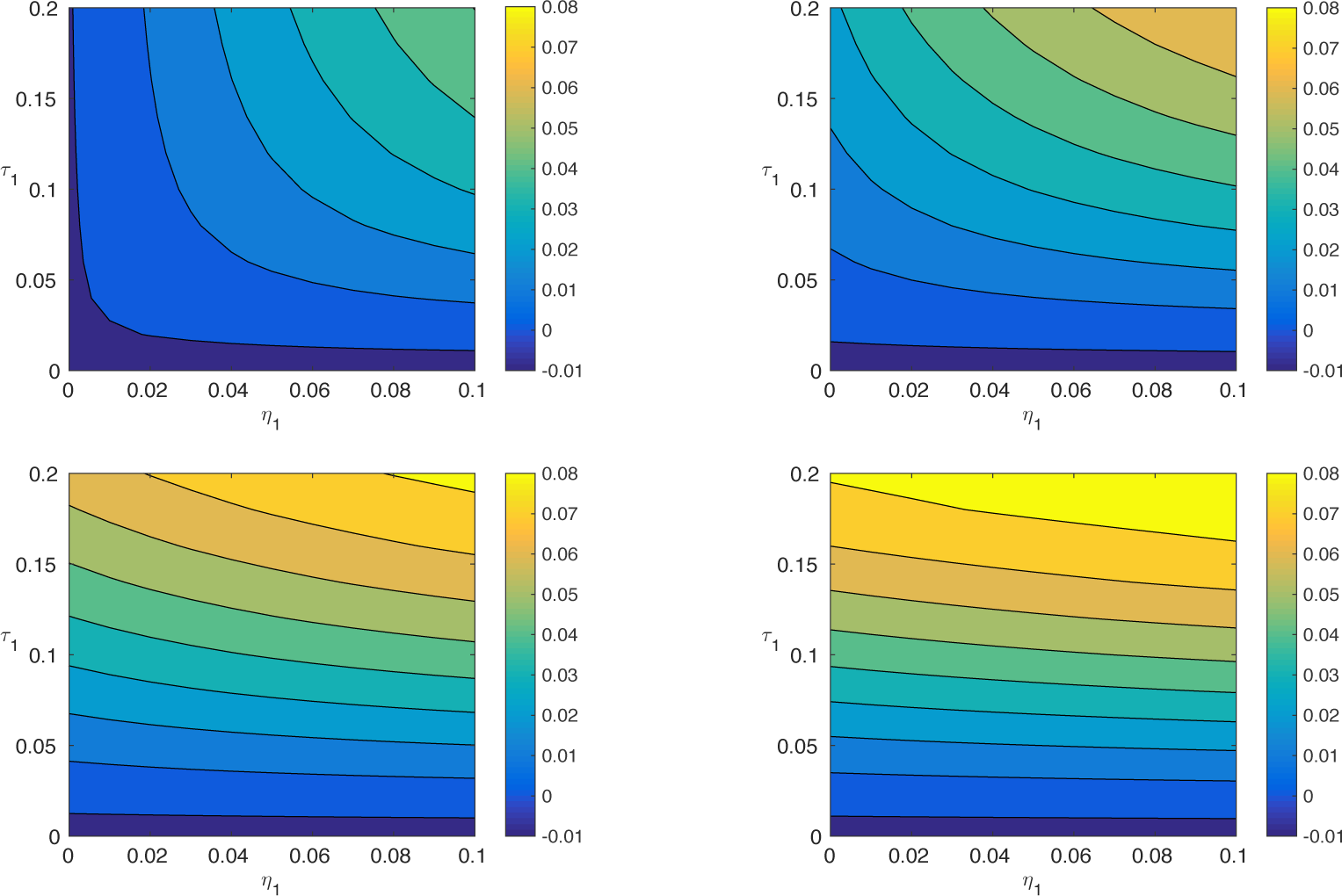
Comparison of early growth with different levels of concurrency: Contour plots of *ΔI*/*I* once the dynamics have entered the early exponential phase. Although eventually the change saturates with larger *k*, it does not saturate nearly as quickly as does the equilibrium size.

This makes sense: if a disease invades a population with serial monogamy, then no matter how infectious it is, it must wait until a partnership ends before it can spread further. However, if there is concurrency, this restriction vanishes, and the spread is not restricted by the partnership time scale. If there are only a few partners, then transmissions lead to localized depletion of susceptibles around an infected individual, stifling further transmissions. As the number of partners increases with a corresponding decrease in transmission rate, the number of transmission opportunities remains the same but the transmissions go to more individuals.

### Caveats

There are a number of caveats of our study that must be highlighted to avoid overinterpreting these results. To simply the model we present, we have neglected many effects. Many of these can be easily incorporated into more sophisticated versions of the model.

- **Acute Phase:** Before mounting an immune response, an individual’s viral load is several orders of magnitude larger than after the immune response develops. During this early phase infectiousness is dramatically increased [20, 21]. If the individual has multiple partnerships, then many more infections can happen in this phase than would be seen if the individual were only in contact with its infector.
- **Heterogeneous degree:** Some people have many more partnerships than others [22]. They generally become infected sooner, and in turn transmit to more individuals. Even if many individuals do not engage in concurrent relationships, if there are some who have many partners, the effects may still be present. This provides an opportunity to reduce disease transmission through an intervention that encourages those without concurrent relationships to select partners without concurrent relationships.
- **Temporal behavior changes:** If the disease dynamics are driven by individuals having periodic high-risk episodes between long-term relationships, then the assumptions of this model are invalid. To correct this, the model must be adapted to allow for periods of high risk behavior, for example when a partnership ends.
- **Age structure:** If there is age-structure in the contact patterns, different effects may be seen. For example, we might think of the younger cohort as a population which has not yet been invaded by infection. In this case, the results about disease invading may be more relevant than the results about population size. Here reducing concurrency could be expected to play an important role in slowing the invasion of this younger cohort.
- **Coital dilution:** We have assumed that the transmission rate scales such that individuals have effectively the same number of sexual acts regardless of their number of partners. This allows us to isolate the effect of concurrency from the effect of frequency of sexual acts. However, if concurrent relationships are associated with more frequent sexual acts, then the conclusions we reach here may not be valid. To correct for this, we need to appropriately weight the transmission rates based on the number of concurrent relationships each partner has.

## Conclusion

We have derived an analytic model which accurately reproduces simulated SI epidemics in a population with concurrent relationships and demographic turnover. We use this model to isolate the role of concurrency in the spread of a disease such as HIV. In isolating concurrency, we consciously choose to neglect a number of other important effects.

Although the model is highly simplistic, it gives insight into the role of concurrency. We see first that the impact of concurrency on the equilibrium size of SI epidemics appears to saturate quite quickly. Consequently we might expect that interventions targeting concurrency will have little impact unless they come close to eliminating concurrent relationships.

However, we see a much larger role for concurrency in determining the early growth rate. As concurrency increases, the early growth is increased, and the impact of concurrency saturates much slower than for the epidemic size. So for slowing the initial invasion of an epidemic, reducing concurrency is likely to have an important impact.

Our observations that the impact of concurrency saturates suggests care may be needed before diverting resources from some intervention to an intervention aimed at reducing concurrency. A more realistic model is needed, and some consideration will need to be made for whether the population is already in equilibrium or whether the epidemic is still growing.

The model presented here is intended as a framework for developing more detailed models. Our goal in introducing this model has been to provide this framework and clearly demonstrate that it is possible to use analytic models to explore disease spread in populations with concurrent relationships with demographic turnover. The predictions our model has provided are true for the simplistic assumptions made. More careful models will be needed to identify conditions under which interventions targeting concurrency will be effective. These models will need to incorporate additional effects such as the acute phase of infection and more realistic information about degree distributions and correlations.

## Supporting Information

Our primary goal in the Supporting Information is to derive the governing equations. For simplicity, we have ignored many effects. Although in the main text we assume all individuals have the same number of concurrent partners, in our derivation here we allow different individuals within the same population to have a different number of concurrent partners as long as all partnerships have the same transmission probability and typical duration. These assumptions could be modified, and a number of other complexities added to the model we develop here, but we do not attempt this now.

### Stochastic population and disease model

We now describe the stochastic rules we assume govern the population and disease dynamics. We use a discrete-time model. We begin with the population dynamics in the absence of disease. At each time step, *N μ* individuals enter the population, and each individual has probability *μ* to independently leave the population. This leads to an equilibrium population size of *N*, but with variation around this value.

Each individual *u* has a constant number of partners *k_u_* which is assigned independently to *u* when *u* enters the population. *P* (*k*) gives the probability that *k_u_* = *k*. We think of *u* as having *ku “*stubs” (also called “binding sites” by [18, 19]). The stubs pair with stubs from other individuals to form partnerships. When a partnership ends the two newly freed stubs join with other free stubs to form new partnerships. We assume that individuals immediately replace their partners so that at the start of each time step all individuals have a full set of partners.^2^

We are interested in the epidemic timescale, which is longer than the individual’s active period. So we must include “birth” and “death” or equivalently immigration and emigration.

We begin with a fraction *ρ* of the population randomly infected. In each time step, multiple events can happen. Since the order of events can matter (a partnership cannot transmit after it ends), we provide a consistent order, shown in figure 3. First, infected individuals transmit to their susceptible partners independently with probability *τ*. Second, individuals may “die” (or leave the population) independently with probability *μ*. Third, *μN* new individuals are added to the population (so the average number present is *N*) and assigned stubs. Fourth, each remaining partnership breaks with probability *η*. Finally, the unpaired stubs form new partnerships, subject to the constraint that old partnerships are not reformed and individuals do not join to themselves. In simulations, these constraints are occasionally not satisfied, in which case the corresponding individuals wait a time step before attempting to reform edges. In a large population, the impact of this failure is negligible, and for our analytic equations below, we can assume that they are satisfied.

## Equation Derivation

### Preliminaries

It will be useful to define the function

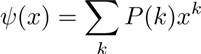

to be the probability generating function of the degree distribution. It has some important properties: *ψ*(1) = ∑*P*(*k*)1^*k*^ = 1, *ψ′*(1) = ∑*kP*(*k*)1^*k*−1^ = 〈*K*〉 where 〈*X*〉 denotes the mean of the random variable *x*.

We now provide a derivation of the deterministic equations governing the large population limit of our model. The derivation is based on [16]. We review the concept of a “test individual” (effectively equivalent to the *cavity state* of [23]). We start with the assumption that the population-scale dynamics are deterministic in the large population limit. A direct consequence of this assumption is the observation that the probability a randomly selected individual has a given status equals the proportion of the population with that status.

Calculating the probability a random individual has a given status turns out to be simpler than calculating the number of individuals in each state. An important observation is that the probability a single randomly chosen individual *u* has a given status is not affected if we prevent it from infecting any other individuals^3^ If we prevent *u* from transmitting to its neighbors, then the status of its neighbors become independent of one another, but this does not alter the probability that *u* is susceptible.

We define a *test individual* to be an individual *u* chosen uniformly at random from the population and prevented from transmitting infection. We have the following sequence of questions which have identical answers if the dynamics are deterministic: Given the initial conditions,

1. What fraction of individuals are susceptible or infected at time *t*?
2. What is the probability a random individual is susceptible or infected at time *t*?
3. What is the probability a randomly chosen test individual is susceptible or infected at time *t*?

In our derivation we assume that a fraction *ρ* is randomly infected at *t* = 0. As long as *ρN* ≫ 1, our equations are appropriate.

To begin our calculations we start with Θ(*t*, *a_u_*), the probability that a stub belonging to an age *a_u_* test individual *u* has never been involved in a transmission to *u*. Once we know that, then the probability a test individual of age *a_u_* and *k_u_* partners is susceptible at time *t* is Θ(*t*, *a_u_*)^*k_u_*^. Averaging this over the entire population of age *a_u_* individuals the probability an age *a_u_* individual is susceptible is

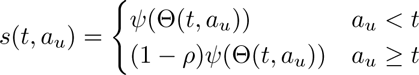

where we recall ∑_*k*_ *P*(*k*)*x^k^* = *ψ*(*x*), and the 1 − *ρ* factor in the second term accounts for the fact that the individual would be infected at *t* = 0 with probability *ρ*.

The probability that a random individual present at time *t* has age *a_u_* is *μ*(1 − *μ*)*a_u_*. The fraction susceptible is thus

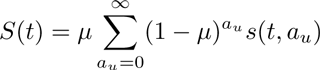

The probability of being infected is

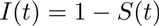

The focus of our calculations is on determining Θ(*t*, *a_u_*). As a boundary condition we have

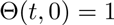

stating that when an individual is first introduced, it has not yet received any infection. Similarly we have the initial condition

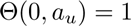

as well, stating that prior to the disease introduction, no transmissions have occurred. The change in Θ in a time step is − *τ*Φ*_I_* where Φ_*I*_ is the probability that the stub has not previously brought infection to *u* and connects to an infected partner at the start of the time step. So we have

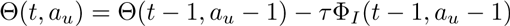

However to do this calculation we require Φ_*I*_ (*t*, *a_u_*) which is still unknown. We can shift our unknown from Φ_*I*_ to Φ_*S*_ (the probability the stub has not transmitted to *u* and currently connects to a susceptible partner) by using

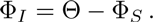

As in calculating *S*, to calculate Φ_*S*_, we turn it into a sum. The probability that a partnership created when *u* joined still exists is (1 *- pb*)^*a_u_*^. The probability that a partnership has some smaller age *a_e_* is *p_b_*(1 − *p_b_*)^*a_e_*^. So

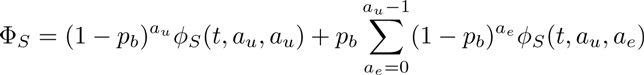

where *ϕ_S_*(*t*, *a_u_*, *a_e_*) is the probability that a stub belonging to an age *a_u_* individual that is part of an age *a_e_* edge has not transmitted by time *t* and *p_b_* is the probability that a stub is freed to find a new partnership (either by death of the partner, or termination of the partnership). The one term outside the sum represents the fact that when the individual first enters the population the stub definitely forms a partnership.

We now find *ϕ_s_*(*t*, *a_u_*, *a_e_*). If the edge formed when *u* was born (*a_e_* = *a_u_*) then this is simply the probability the partner *v* is susceptible given that *v* has an age *a_u_* partnership with *u*, which we denote *χ*(*t*, *a_u_*). However, if the edge formed after *u* was born (*a_e_* < *a_u_*), then *ϕ_S_* (*t*, *a_u_*, *a_e_*) is the probability Θ(*t* − *a_e_*, *a_u_* − *a_e_*) that the stub was not responsible for transmitting infection to individual *u* prior to the current partnership forming times *χ*(*t*, *a_e_*). As Θ(*t* − *a_u_*, 0) = 1 these coincide when *a_u_* = *a_e_*, so we can write

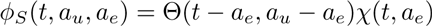

We now find χ(*t*, *a_e_*) similarly to *s*(*t*, *a_u_*). It is

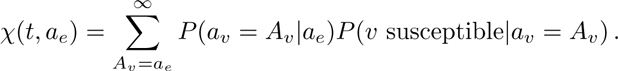

If *a_e_* ≥ *t*, then we know that *v* was born either when the disease was introduced or earlier. Thus no previous partnership could have transmitted to *v*. if we assume *a_v_* = *A_v_* ≥ *a_e_*, then the probability *v* is susceptible is the probability that it escaped infection when the disease was introduced 1 − *ρ* times the probability that it has not been infected by any other partners. Because of how *v* is selected (it is *u*’s partner), *v* is likely to have a higher degree than a randomly selected individual. The probability *v* has deg ree *k_v_* = *k* is *kP*(*k*)/ 〉*K*〈. So the probability *v* is susceptible given *A_v_* is (1 − *ρ*)∑_*k*_[*kP*(*k*)/〈K〉]Θ(*t*, *A_v_*)^k−1^ = (1 − *ρ*)*ψ′*(Θ(*t*, *A_v_*))/〈K〉. Thus for *a_e_* ≥ *t* we have

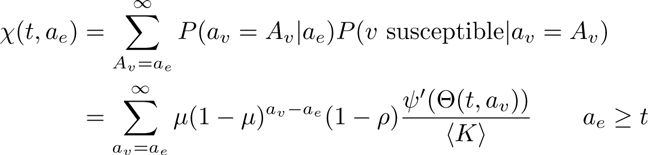

For *a_e_* < *t* there are three important cases to consider based on whether the partner was born before or at the same time that the partnership was formed and whether the partner was born before or after the disease was introduced.

- If the edge formed when *v* was born then *A_v_* = *a_e_* for which *P*(*a_v_* = *a_e_*|*a_e_*) = (1 − *P_e_*) and *P*(*v* susceptible|*a_v_* = *a_e_*) = ∑_*k*_[*kP*(*k*)/〈*K*〉]Θ(*t*, *a_e_*)^*k*−1^ = ψ′Θ(*t*, *a_e_*)/〈*K*〉, which measures the probability that another partner of *v* has not transmitted to *v*.
- If *v* was born before the edge formed but after the disease was introduced then *a_e_* < *A_v_* < *t* and *P*(*a_v_* = *A_v_*|*a_e_*) = *P_e_μ*(1 − *μ*)^*A_v_* − *a_e_* − 1^. Although *u* has not transmitted to *v*, it is possible that a previous partner of *v* that was eventually replaced by *u* did. Thus the probability *v* is susceptible is Θ(*t* − *a_e_*, *A_v_* − *a_e_*)*ψ′*(Θ(*t*, *A_v_*)).
- If *v* was born before the disease was introduced, then *A_v_* ≥ *t*. We again have *P*(*a_v_* = *A_v_*|*a_e_*) = *P_e_μ*(1 − *μ*)^*A_v_* − *a_e_* − 1^, but there is an extra factor of 1 − *ρ* in the probability *v* is susceptible. *P*(*v* susceptible|*A_v_*) = (1 − *ρ*)Θ(*t* − *a_e_*, *A_v_* − *a_e_*)*ψ′*(Θ(*t*, *A_v_*)).

So for *a_e_* < *t* we have

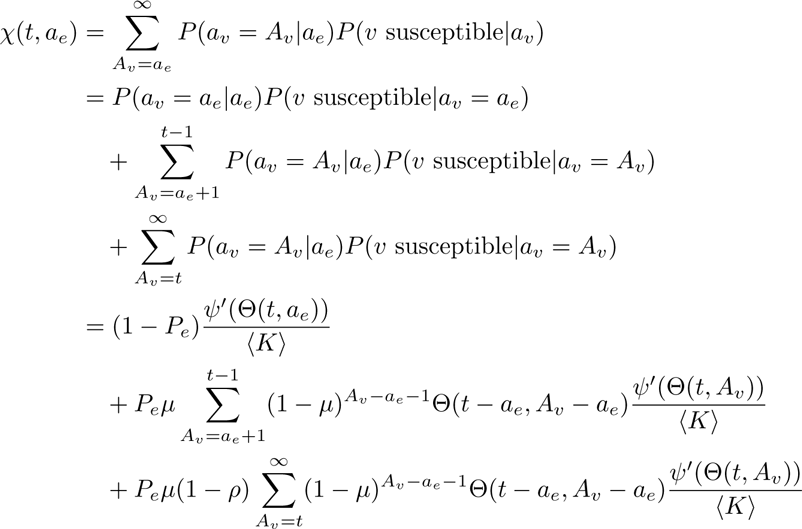

### Simplification for *a_u_* > *t*

We claim that the value of Θ(*t*, *a_u_*) is the same for all *a_u_* ≥ *t*. This follows from the fact that at *t* = 0 all the values are 1. By inspecting the equations for the evolution of Θ, we see that if we assume Θ(*t*, *a_u_*) is the same for all *a_u_* ≥ *t*, then the change in Θ is also the same. Thus we can assume Θ(*t*, *a_u_*) = Θ(*t*, *t*) if *a_u_* > *t*. This argument would break down if partnership formation were affected by age differences.

Among the resulting simplifications is the observation that for *a_e_* ≥ *t*, the expression for *χ*(*t*, *a_e_*) simplifies to (1 − *ρ*)*ψ′*(Θ(*t*, *t*))/〈*K*〉.

### Governing Equations

Our full system of equations becomes

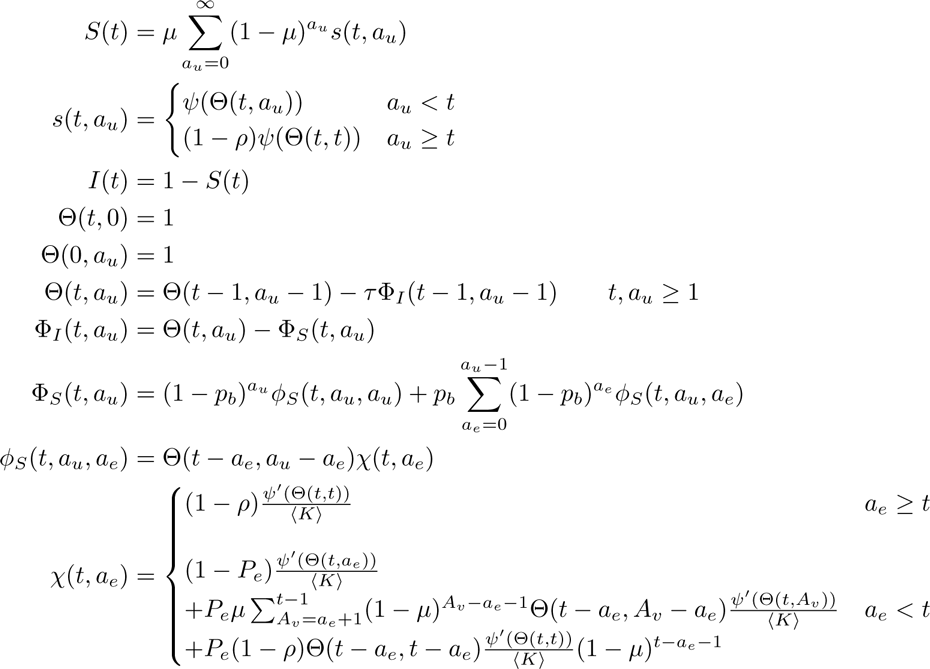

We can derive a differential equations version of this by replacing the time step *n* with the time step *nΔt*, and then calculating derivatives in the *Δt* → 0 limit.

A differential equations version of this can be derived by treating the time step as *Δt* rather than 1 and assuming that the event probabilities are all proportional to *Δt*. Then taking *Δt* → 0 yields differential equations. We will use 
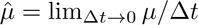
 and similarly define other variables.

We find

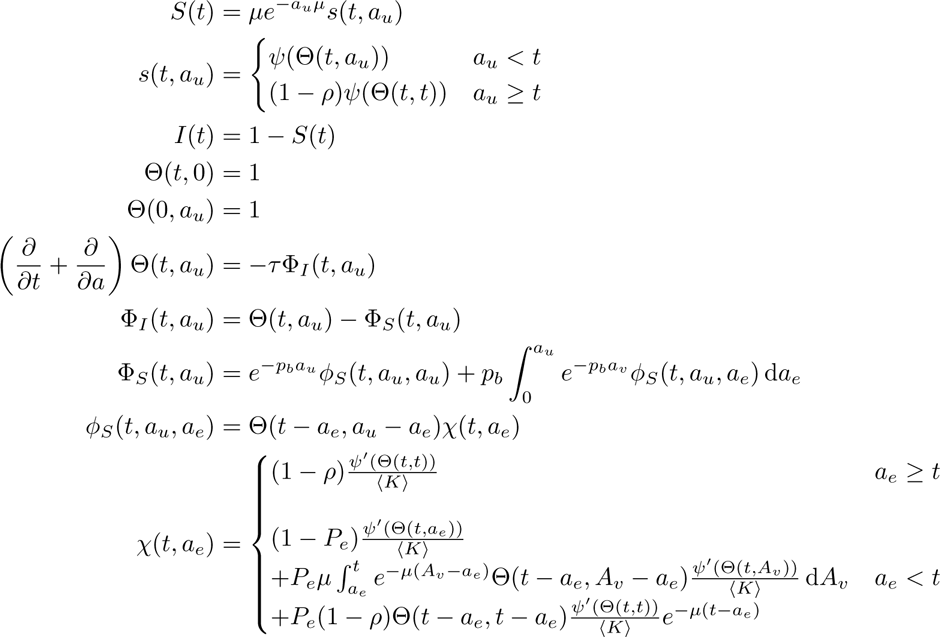

The simplest numerical method to solve this system of equations would apply an Euler method, which corresponds to solving the discrete-time equations above.

## Footnotes

1 However, the probability of a Bobbie to Alex to Charlie transmission chain is slightly reduced because some of the interactions between Alex and Charlie will have already happened by the time Alex is infected.

2 In [16] there is discussion of how to include more complicated partnership dynamics.

3 Although it is not necessary here, it may be helpful to recognize that that the assumption the stochastic process exhibits deterministic population-scale dynamics means that a change of out come for a vanishingly small fraction of events does not alter the population-scale dynamics.

